# Development and Application of a Novel Simple Sequence Repeat Mining Algorithm Based on Regular Expression

**DOI:** 10.1101/2022.06.01.494292

**Authors:** Zhenguo Jia, Ruimei Geng, Xiuming Wu, Shuai Chen, Ying Tong, Aiguo Yang, Chenggang Luo, Min Ren

## Abstract

Simple sequence repeats (SSRs) are molecular genetic markers that are powerful tools in genomics studies; SSR markers are routinely mined as a part of genetic workflows. Here, we developed a novel SSR mining algorithm based on regular expression that can reduce the complexity of commonly used SSR mining software. We used the following SSR mining regular expression: ({i, j}?) (\1) {k}, where i and j denote the minimum and maximum lengths of the motifs of the SSR sequence, respectively, and k is the minimum number of repeat motifs. From this SSR mining algorithm, we developed an SSR sequence analysis software (named “regexSSRw”) that is capable of mining eligible SSR loci from FASTA format sequences; regexSSRw can be accessed at https://github.com/renm79/rgxSSRw. This SSR mining algorithm can aid a range of applications, from being used by programmers in the development of SSR mining software to being implemented by scholars into their SSR marker workflow.

## Introduction

Simple sequence repeats (SSRs) refer to tandem repeats of 1–6 nucleotides in deoxyribonucleic acid (DNA) sequences. Many SSRs are widely interspersed within the genomes of eukaryotes and most prokaryotes. SSRs are both abundant in polymorphisms and stable, which allows for their transmittance to offspring, according to Mendelian inheritance. SSRs are ideal molecular markers because they can discriminate between homozygous and heterozygous genotypes, are co-dominantly inherited, possess abundant variation, are randomly distributed throughout the genome, can be rapidly identified and analyzed by polymerase chain reaction (PCR), and require relatively low amounts of DNA for high quality results. Numerous primer sequences have been published, with high repeatability being a common feature. Consequently, SSR technology has been widely used in genetic studies into plants and animals (Vieira et al.,2016;Rajesh et al., 2021;Malhotra et al., 2021;Yang et al., 2021). SSR sequence analysis and mining has thus become a fundamental process in structural genomics research.

Currently, SSR mining algorithms and widely used software such as Tandem repeats finder(Benson,1999), SSRIT (Temnykh, 2001), TROLL (Castelo et al., 2002), MISA (Thiel et al., 2003),SSR primer(Robinson et al.,2004), MSATCOMMANDER (Faircloth, 2008), FullSSR(Metz et al.,2016) and SSRminer (https://github.com/CJ-Chen/TBtools-Manual), are mostly based on string searches. However, some features of SSRs produce large numbers of redundant search results in these algorithms. For instance, a particular repetitive SSR sequence such as “AGAGAGAGAG” will be subjected to repeated statistics, as multiple search strings are used (“AG,” “AGAG,” “AGAGAG,” etc.); the sequence “CTCTCTCT” will also be subjected to repeated statistics with the search strings of “CT” and “TC.” Thus, it is important to remove redundant results using statistical or logical analysis strategies. However, this greatly increases the complexity of the algorithms and code required to develop relevant programs. In particular, this causes difficulties and inconveniences regarding the quick preparation of scripts and high-throughput sequence analyses. To address this, here, we explored a novel algorithm that utilizes regular expression to mine SSRs and avoid redundant searches, with the aim of significantly simplifying the structure of the program to improve execution efficiency, increase maintainability, and enhance readability.

## Methods

### Algorithm Flow

We used the computer language Python 2.7.2 (which can be downloaded from https://www.python.org/downloads/release/python-2713/) to create the algorithm used in this study. Our novel algorithm includes several steps regarding its use in the SSR mining workflow (Fig. 1). First, the calculation parameters should be determined based on the purpose of the experiment; the minimum and maximum lengths of motifs and the number of repeat motifs for the SSR sequence being mined should also be determined. Second, a regular expression that is capable of specifically matching SSR sequences should be constructed. This expression should then be used to analyze the target sequence, where no matches indicate the absence of any SSR loci, whereas sequence matches pinpoint usable SSR loci. In general, this algorithm must be able to determine whether a mined SSR locus comprises single bases; these need to be removed such that the final SSR loci can be obtained.

**Fig. 1.**
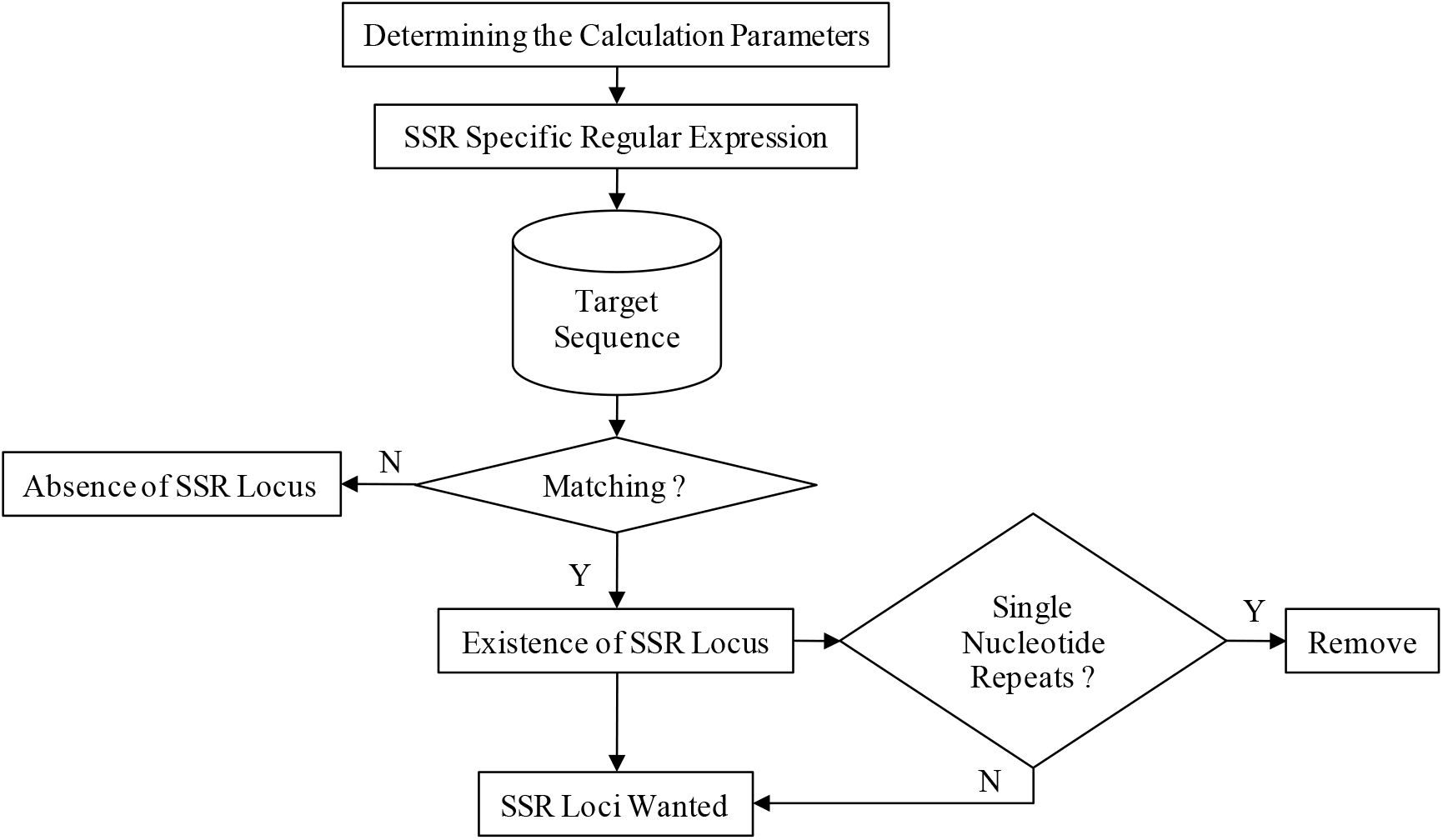
Workflow for SSR mining algorithm developed in this study

### Regular Expression Specific Matching SSR Sequence

The base compositions of SSR sequences exhibit distinctive characteristics. Specifically, they contain repeated motifs that usually consist of one to eight bases; they are formed by the random combination of the four bases: A (adenine), T (thymine), C (cytosine), and G (guanine). Owing to the sequence characteristics of SSR, we constructed the following specific matching SSR regular expression:

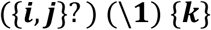

where i and j denote the minimum and maximum lengths of the SSR sequence motifs, respectively, and k is the minimum number of repeat motifs. For example, when the algorithm determines that it needs to mine an SSR locus with a motif length of 2–6 bases and requires at least five repeats, the regular expression should be ({2,6}?) (\ 1) {4}.

## Results

### Algorithm Implementation

In this study, we applied the Python computer language to program an SSR sequence mining class based on the aforementioned algorithm (https://github.com/renm79/rgxSSRw/blob/main/SSR_class.py). This SSR sequence mining class can be easily used during the preparation of SSR mining software. When using this mining algorithm, the target sequence, the minimum and maximum lengths of the motif, and the number of repeats all needed to be defined; the default values were an SSR locus with a minimum length of 2, a maximum length of 6, and a minimum of five repeats. The SSR sequence mining class had six output parameters. CountSSR denotes the number of SSR loci returning to the sequence to be analyzed, with a value of 0 indicating the absence of SSR loci. ListSSR commands the algorithm to return all SSR loci in the form of a list, whereas ListMotif lists all the motifs. GetLeft and GetRight return the upstream and downstream sequences of an SSR locus, respectively. GetPosition indicates the position of a certain SSR locus in the sequence to be analyzed.

### Development of SSR Mining Procedures

Based on the above SSR sequence mining class, we developed an SSR sequence analysis software named “regexSSRw” (Fig. 2). It was programmed using the Python computer language, which can be run on Windows, Linux, MacOS, and other operating systems. The software is capable of mining eligible SSR loci from FASTA format sequences, in accordance with preset parameters. Currently, this software is available via the Github hosting service; it can be accessed at https://github.com/renm79/rgxSSRw.

**Fig. 2.**
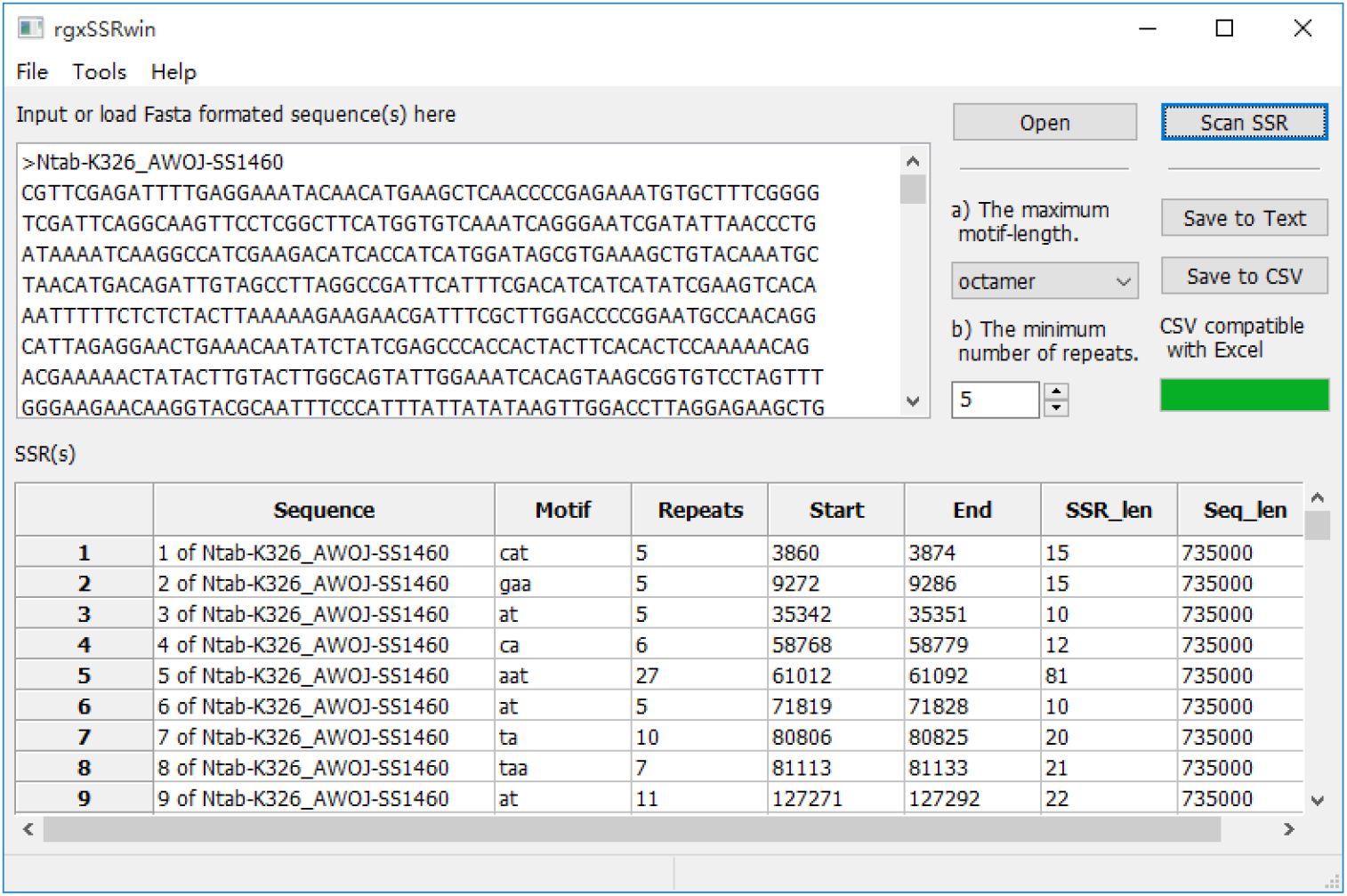
User interface for SSR analysis software rgxSSRwin.

### Algorithm Validation

To verify the validity of the algorithm, we downloaded a scaffold sequence with 735,000 nucleotide bases, Ntab-K326_AWOJ-SS1460.fasta (Sierro et al.,2014), which we randomly selected from a *Nicotiana tabacum* reference genome acquired from the Sol Genomics Network website. We used rgxSSRwin and two widely used SSR mining software programs, Sputnik and SSRfinder, to screen the distribution of SSR loci in the *N. tabacum* sequence, using the same search parameters (minimum SSR motif length = 2, maximum SSR motif length = 8, and minimum motif = 5). The results showed that rgxSSRwin and SSRfinder both found 100 SSR loci in the same locations but that Sputnik only found 75 SSR loci. Based on these results, we determined that the regular expression-based SSR mining algorithm used in this study is both accurate and reliable.

## Discussion

Today, SSR marker research has become an important aspect of molecular research, and has allowed for increased technological developments that are now widely used in human, animal, and plant genomic studies. For example, many important crop species, such as bread wheat, barley, rice, and soybean, have large numbers of available SSR markers (Zheng et al., 2020; Qi et al., 2021; Chen et al., 2020; Zhang et al., 2021).However, many species genomes have not been explored for SSR markers; they are in need of SSR mining. Fortunately, publicly available sequences for SSR mining are increasingly being published online, allowing for easy access. Scripting languages such as Perl, Python and R are often utilized to collate and analyze vast amounts of sequence information in bioinformatics research. As SSR sequence mining and marker development have become routine tasks in DNA sequence analysis, it is important to develop efficient workflows for this process. However, SSR mining is limited by the complexity of its algorithms; they increase the complexity of script programming, indicating that researchers with high programming skills are required to carry out SSR mining functions via scripts. When utilizing conventional SSR analysis software, these complex algorithms require increased operational processes to handle the data analysis and the large number of search results; this reduces work efficiency. Thus, here, we proposed a novel SSR mining algorithm that utilizes regular expression to resolve redundancy issues in analysis results by reducing the complexity of the algorithm. Reducing the complexity during SSR mining significantly improves the analysis efficiency of DNA sequences, because DNA sequences are essentially a form of text string; regular expression is a powerful tool for manipulating text strings. Thus, in this study, we demonstrated the convenience and efficiency of using regular expression to analyze DNA sequences; we predict that it could be widely applied and used in future genomic work.

## Conclusion

In this study, we developed a new SSR mining algorithm that solves the problem of redundancy of analysis results by using regular expressions, thus reducing the complexity of the algorithm. This facilitated the implementation of an SSR mining function in script programs, and significantly improved the efficiency of DNA sequence analysis.

